# Optimal transcriptional regulation of cellular responses to sudden environmental shifts

**DOI:** 10.1101/2020.09.14.297317

**Authors:** Daniel Schultz, Lev S. Tsimring

## Abstract

Cellular responses to sudden changes in their environment require prompt expression of the correct levels of the appropriate enzymes. These enzymes are typically regulated by transcription factors that sense the presence of inducers and control gene expression for the duration of the response. The specific choice of regulatory strategy depends on the characteristics of each cell response, with the pattern of gene expression dictated by parameters such as the affinity of the transcription factor to its binding sites and the strength of the promoters it regulates. Although much is known about how gene regulation determines the dynamics of cell responses, we still lack a framework to understand how the many different regulatory strategies evolved in natural systems relate to the constraints imposed by the selective pressures acting in each particular case. Here, we analyze a dynamical model of a cell response where expression of a transcriptionally repressed enzyme is induced by a sudden exposure to its substrate. We identify strategies of gene regulation that optimize the response for different types of selective pressures, which we define as a set of costs associated with substrate, enzyme and repressor intracellular concentrations during the response. We find that regulated responses happen within a defined region in the parameter space. While responses to costly (toxic) substrates favor the usage of strongly self-regulated repressors, responses where expression of enzyme is more costly than its substrate favor the usage of constitutively expressed repressors. There is only a very narrow range of selective pressures that would favor weakly self-regulated repressors. This framework can be used to infer which costs and benefits are most critical in the evolution of natural examples of cellular responses, and to predict how a response can optimize its regulation when transported to a new environment with different demands.

## INTRODUCTION

Cellular responses to changes in their environment, in fact cellular processes in general, require the coordinated expression of genes. When exposed to a chemical compound that demands urgent processing, the cell needs to sense its presence and synthesize precise amounts of correct enzymes in a timely fashion. Therefore, the optimization of responses does not involve solely the evolution of enzymes, but also the evolution of appropriate regulation that can manage the costs and benefits associated with the expression of these enzymes over the course of the response [1]. Both of these aspects evolve in concert in the cell’s DNA sequence: while the coding regions (genes) contain information about the biochemical properties of enzymes and regulators, the non-coding regions contain information about DNA binding sites that together dictate the patterns of enzyme expression. Much attention has been given to the evolution of genes [2], while other studies recognize that the evolution of cell responses typically involve changes in regulatory pathways [3, 4]. However, we still lack a framework to describe the evolution of cellular responses as dynamical processes [5–8] that can be directly inferred from genotype.

The balance of costs and benefits shaping the evolution of genes and their regulation depends on the particular context of each cell response [9]. The enzymes at play invariably interfere with other cell processes, so their expression often poses tradeoffs between their intended function and their unintended consequences. For instance, the presence of a toxin is harmful for the cell, but overproduction of the enzyme that rids the cell of this toxin can disrupt other cell functions, posing heavy metabolic burdens and impairing cell growth [10]. In this case, the choice of regulation of such enzyme will depend strongly on the relation between the cost of the presence of the toxin versus the cost of enzyme side effects. Ultimately, the selective pressures guiding the evolution of a cell response can be translated into a set of costs associated with the time-dependent concentration of each of the molecular components participating in the response. Evolution then proceeds to minimize the costs, by optimizing the rates of the biochemical interactions involving these components.

Here, we analyze how selective pressures shape the evolution of a typical cell response pathway: a transcription factor regulating an enzyme for a particular substrate (Fig. 1A). In the absence of the substrate, the expression of the enzyme is repressed by the transcription factor. In the presence of the substrate, however, as its concentration starts accumulating in the cytoplasm, the transcription factor recognizes and binds the substrate, losing affinity to the DNA and de-repressing the enzyme. The enzyme then proceeds to process the substrate, until its intracellular levels begin to decrease. Finally, when the intracellular concentration of the substrate returns to low levels, the transcription factor is released and resumes repression of the enzyme. We focus on the case of a sudden shift in the substrate concentration eliciting a fast response, bringing the system out of equilibrium, such as in the case of a bacterium responding to an environmental stress.

**FIG. 1:**
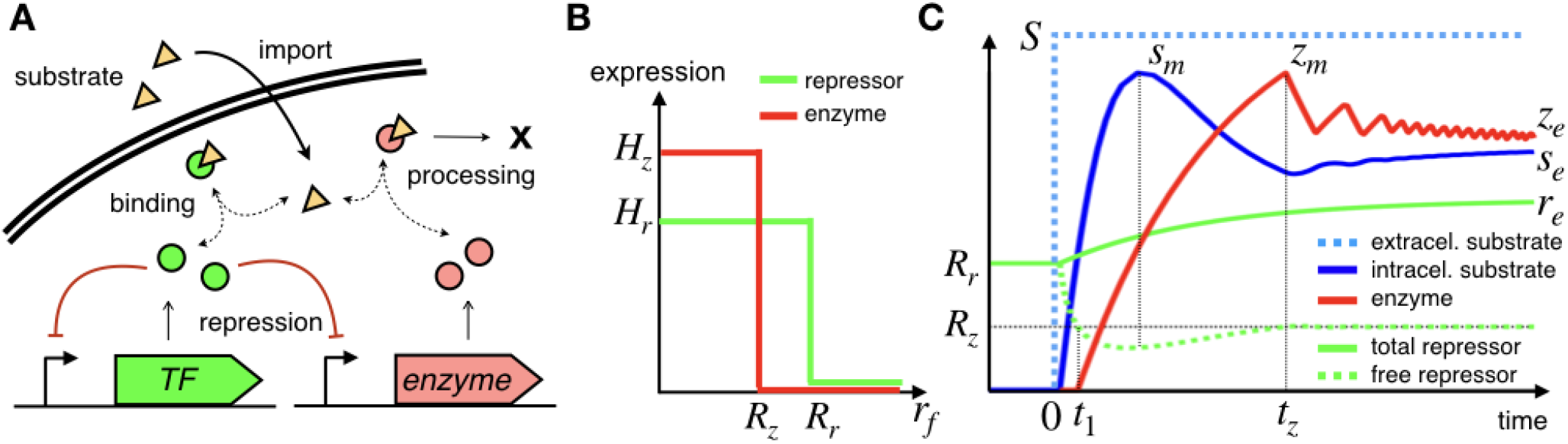
General model of cell responses. **A** In the absence of substrate, a transcription factor represses the expression of an enzyme and of itself. When substrate is imported into the cell, it binds and inactivates the repressor, releasing expression of both genes, and the enzyme proceeds to process the substrate. Cell components are degraded or diluted due to cell growth. **B** Regulation of repressor and enzyme are modeled as “on/off” according to the concentration of free repressor. Repressor and enzyme are fully expressed at rates *H_r_* and *H_z_* when free repressor levels are below thresholds *R_r_* and *R_z_*, respectively, and are not expressed if free repressor concentration exceeds these thresholds. **C** A typical response to a sudden increase in extracellular substrate concentration *S* at time zero, showing subsequent peaks in intracellular substrate (*s_m_*) and enzyme (*z_m_*) concentrations, before stabilizing to final values *s_e_*, *z_e_* and *r_e_*.

## GENERAL MODEL OF CELL RESPONSES

We model the cell response by a system of three differential equations describing the concentrations of the enzyme *z*, the repressor *r* and the substrate *s* [11]:

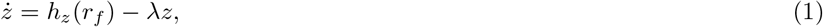

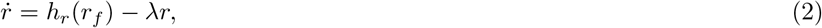

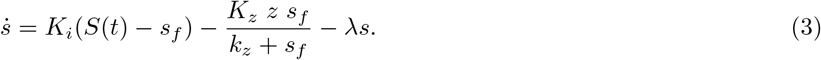

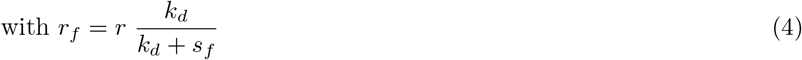

The substrate is imported into the cell with a rate proportional to the difference between its extracellular (*S*) and intracellular (*s*) concentrations with an import rate constant *K_i_*, as is the case when the substrate diffuses across the cell membrane. The substrate is processed by the enzyme *z* following Michaelis-Menten kinetics, with a catalytic rate constant *K_z_* and a Michaelis constant *k_z_*. As intracellular substrate binds and inactivates the repressor, the synthesis of the enzyme and the repressor inside the cell is regulated by the amount of free (not bound to the substrate) repressor *r_f_* according to regulatory functions *h_z_* and *h_r_*, respectively (discussed below). All intracellular components are diluted in the cytoplasm as the cell grows with a rate λ, which we consider constant for the duration of the response. Since the biochemical binding and unbinding of the substrate to the transcription factor typically happens at a much faster rate than the aforementioned processes, we consider their unbound (free) forms (*s_f_, r_f_*) to be in chemical equilibrium with the bound form [rs] with a dissociation constant *k_d_*, such that *r_f_s_f_* = *k_d_*[*rs*]. Therefore, the inactivation of repressor by the substrate is included in the differential equations implicitly, with *r_f_* calculated from this binding equilibrium.

We consider a simple “on/off” regulation of the synthesis rates of enzyme and repressor (*h_z_*, *h_r_*) by the concentration of free repressor *r_f_* (Fig. 1B). Synthesis is kept at a constant rate (*H_z_*, *H_r_*) at low levels of free repressor and is shut down when the concentration of free repressor crosses a threshold for inhibition (*R_z_*, *R_r_*). This description results in four parameters describing gene regulation during the response, corresponding to promoter strengths and thresholds of inhibition of each the enzyme and its repressor. Although the “on/off” description misses finer details of the regulation [12, 13] that are important in setting the steady-state concentrations of the molecular components of the response, it still captures essential aspects of the dynamics of cell processes out of equilibrium. “on/off” gene expression implements “bang bang” control, ubiquitous among cell processes, which is frequently the optimal solution for minimum-time problems [14]. For instance, the quickest way to reach a specific concentration of enzyme is to synthesize it at maximum speed until a switching point where expression is shut off. “on/off” expression also captures the temporal organization of gene expression in pulses, typical of cell processes such as stress responses, signaling and development [5, 15–18].

We can now numerically evaluate the time course of all the relevant concentrations in a typical response where the extracellular concentration of substrate suddenly rises from zero to a high level *S* at time zero (Fig. 1C). In the absence of substrate, the concentration of repressor will sit at the threshold for its own repression *R_r_* (provided *H_r_*/λ > *R_r_*). Decreases in repressor concentration will activate its synthesis, while increases in its concentration will shut it off. Provided that *R_z_* < *R_r_*, the enzyme concentration will be zero. Once the substrate is present, it starts to flow into the cell and bind the repressor, decreasing the amount of free repressor until it crosses the threshold for activation of enzyme *R_z_* at time *t*_1_. The enzyme is then expressed, processing substrate until its intracellular concentration starts to decrease, after reaching a peak concentration *s_m_*. The concentration of free repressor then starts to increase until it crosses back the threshold for enzyme repression *R_z_* at time *t_z_*, after which the concentration of enzyme begins to decrease. After a series of decaying oscillations in substrate and enzyme concentrations, caused by filtering delays in the regulatory negative feedback loop, during which the enzyme synthesis is intermittently switched on and off, the levels of response components equilibrate to their final steady-state values. In this process, free repressor concentration equilibrates to *R_z_*, while the repressor synthesis is unchecked, reaching maximum levels.

We further consider two approximations. Inequality *s_f_ ≪ k_z_* assumes that the concentration of intracellular substrate does not reach high enough levels to saturate the enzyme. This regime is typical of fast responses, which are readily induced by the substrate and are pressured to process it rapidly. Inequality *r_f_ ≪ k_d_* assumes that the concentration of intracellular substrate is much higher than the concentration of repressor, such that *s_f_ ≈ s*. This means that regulation operates at relatively low levels of repressor, as is usually the case, and repressor binding does not significantly reduce the concentration of free substrate. We also rescale the variables and parameters of the system, as described in the SI and summarized in table I, to arrive at non-dimensional equations:

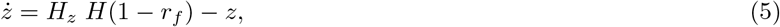

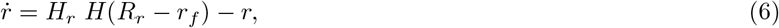

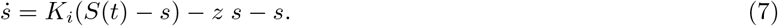

with *r_f_* = *r*/(1 + *s*) and *H*(*x*) the Heaviside function. In these rescaled variables without loss of generality we choose the dilution rate λ and the threshold repressor value for activator *R_z_* to be unity. The regulation of the system is now described by three non-dimensional parameters. 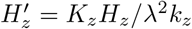 is the enzyme level at full expression, with enzyme levels rescaled to the enzyme catalytic rate. 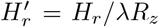 is the repressor level at full expression, with repressor levels rescaled to the threshold for enzyme repression. 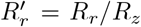 is the threshold of repressor self-repression in relation to the threshold for enzyme repression (primes are dropped below for convenience).

**TABLE I:** Description of variables and parameters used in the model (primes are dropped in the main text for simplicity)

| Variables |  |
| --- | --- |
| $t$ | time |
| $z$ | enzyme concentration |
| $r$ | total repressor concentration |
| $s$ | intracellular substrate concentration |
| $S$ | extracellular substrate concentration |
| $r_f$ | free (active) repressor concentration |
| $s_f \approx s$ | free intracellular substrate concentration |
| Parameters |  |
| $\lambda$ | cell growth rate |
| $K_i$ | substrate import rate |
| $K_z$ | enzyme catalytic rate |
| $k_z \gg s_f$ | enzyme Michaelis constant |
| $k_d \gg r_f$ | repressor/substrate dissociation constant |
| $H_z$ | enzyme synthesis rate (when *on*) |
| $H_r$ | repressor synthesis rate (when *on*) |
| $R_z$ | threshold for enzyme repression |
| $R_r$ | threshold for repressor self-repression |
| Dimensionless Variables |  |
| $t'$ | $\lambda t$ |
| $z'$ | $K_z z / \lambda k_z$ |
| $r'$ | $r / R_z$ |
| $s'$ | $s / k_d$ |
| $S'$ | $S / k_d$ |
| $r'_f$ | $r_f / R_z$ |
| Dimensionless Parameters |  |
| $K'_i$ | $K_i / \lambda$ |
| $H'_z$ | $K_z H_z / \lambda^2 k_z$ |
| $H'_r$ | $H_r / \lambda R_z$ |
| $R'_r$ | $R_r / R_z$ |

We can now theoretically analyze the time course of a response to a sudden increase in extracellular levels of substrate. For this, we can divide the time course of the response by the events when the concentration of free repressor crosses the thresholds for repression of either the enzyme or the repressor itself. In the intervals between these events, the synthesis rates of both the enzyme and the repressor are constant, and the model equations can be integrated explicitly. We detail this procedure in the SI. Next, we consider the different regimes of the cell response in the parameter space defined by *H_z_*, *H_r_* and *R_r_*.

## OPTIMIZATION OF RESPONSE DYNAMICS

In this section we will discuss how selective pressures in the cell environment determine the choice of parameter values that optimize the response. Let us fix the enzyme full expression level *H_z_* and only consider the two-parameter plane *R_r_* × *H_r_*, which describes the regulation by the transcriptional repressor (Fig. 2). This plane can be divided into regions where the progressions of the response are qualitatively different, and the boundaries between these different regimens can be used to constrain the search for optimal parameters. *Constitutive enzyme expression:* when *H_r_* < 1 full expression of repressor does not reach the threshold for enzyme inhibition, and when *R_r_* < 1 the repressor represses itself more strongly than it represses the enzyme. In both of these cases, the repressor is unable to regulate the enzyme, which is always expressed at a constant level *H_z_* (Region 0). *Constitutive repressor expression:* when *R_r_* > *H_r_*, the threshold for repressor self-repression is higher than its full-expression level. Since self-repression is never reached, the repressor is always expressed at a constant level *H_r_* (Regions 1, 3 and 5). Notably, constitutive expression of repressor in this regions is independent of *R_r_*, so any point where *R_r_* > *H_r_* is equivalent to the point in the boundary where *R_r_* = *H_r_*, where constitutive repressor expression is already reached. *Repressor levels for regulated response:* at very high levels of repressor expression *H_r_* > *α*_2_ = *K_i_ S*/(*K_i_* + 1) (or very low substrate level *S*), the concentration of free repressor never drops below the threshold to activate the enzyme, so the response is never induced (Regions 5 and 6). At lower levels of repressor expression 1 < *H_r_* < *α*_1_ = *K_i_ S*/(*K_i_* + *H_z_* + 1), repressor levels are insufficient to curb enzyme expression once the response is induced, leading to unnecessary full expression of enzyme (Regions 1 and 2). Therefore, regardless of the selective pressures applied, optimized parameters for the response can be expected to be found in the region defined by 1 < *R_r_* < *H_r_* and *α*_1_ < *H_r_* < *α*_2_ (Region 4), with the remaining parameter space either being equivalent to the boundaries of this region or presenting extreme and obviously suboptimal solutions.

**FIG. 2:**
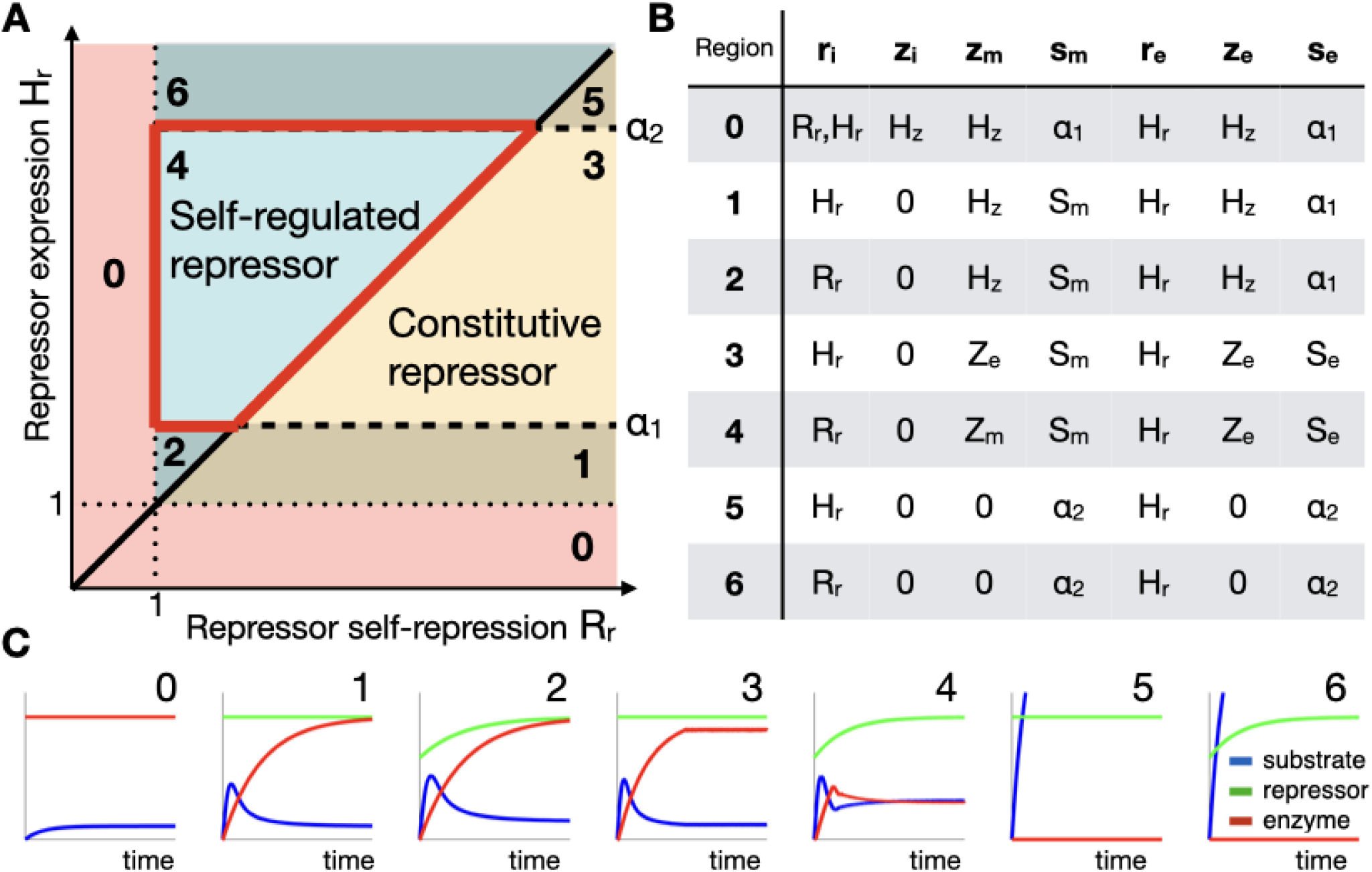
Distinct progressions of cell responses. **A** Regions in the *R_r_* × *H_r_* plane that correspond to qualitatively different response progressions. Region 4, highlighted in red boundaries, contains the parameters resulting in all possible regulated responses, where the enzyme is repressed in the absence of substrate and is induced in its presence. **B** Values of the key concentrations in the time course of the response: *r_i_* and *z_i_* are the initial repressor and enzyme concentrations, *z_m_* and *s_m_* are the peak values of enzyme and substrate, and *r_e_, z_e_* and *s_e_* are the end values of repressor, enzyme and substrate, respectively. Expressions for *α*_1_, *α*_2_, *Z_e_*, *Z_m_, S_e_, S_m_* are summarized in the SI. **C** Time courses of the responses corresponding to the different regions in the *R_r_* × *H_r_* plane in A.

The selective pressures acting on the system during its evolution dictate the choice of parameters that maximize its efficiency. An ideal response should achieve maximal processing of the substrate, while expending the least amounts of enzyme and repressor. Therefore, the optimization of the response involves the minimization of substrate, enzyme and repressor levels during the course of the response. We note that minimization of substrate levels is desirable both in the case of a drug response, where there is a cost associated with the toxicity of the substrate, or in the case of a metabolic pathway, where there is a benefit associated with the processing of the substrate. The specific context of the system operation poses additional constraints that might prioritize the minimization of one of the response components over the other. For instance, the choice between minimization of substrate or enzyme levels depends on which of them imposes higher costs for the cell. In addition, costs associated with each component can depend not only on the maximum concentrations reached, but also on the duration for which they are sustained. Strictly speaking, the system performance is characterized by the whole time courses of substrate, enzyme, and repressor levels during the response, which makes the optimization problem infinite-dimensional. To keep the dimensionality of the problem finite, we approximate the total cost of a given response using a finite set of key concentrations *c_i_*: initial, maximum and final levels for each of the three components (substrate *s_i_, s_m_, s_e_*, enzyme *z_i_*, *z_m_, z_e_*, and repressor *r_i_*, *r_m_, r_e_*). Since we always have *r_m_* = *r_e_*, and we always assume that *s_i_* = 0, we are left with seven output metrics. We then define the selective pressure acting on the system by a set of weights *w_i_* assigned to each of these metrics, and compute the total cost of the response as a weighted sum of individual metrics *C* = ∑_*i*_ *w_i_c_i_*. The relative weights of different metrics determine which set of regulatory parameters (*H_z_,H_r_, R_r_*) minimizes the total cost *C*. This approach assumes for simplicity that the cost grows linearly with each key concentration, which might not hold true, particularly in cases where the cost saturates at specific levels.

Next, we analyze the importance of the cost of individual components in determining the optimal response regulation. The initial levels of components *r_i_* and *z_i_* are sustained in the absence of substrate. Therefore, if the presence of substrate and demand for the enzyme are relatively rare, the cost of these initial levels should be high to reflect a selective pressure to keep the system repressed by default. Otherwise, constitutive expression of appropriate levels of the enzyme is optimal. For regulated responses, if we disregard the costs of expressing repressor and consider only the costs associated with substrate and enzyme, optimal solutions are found at the highest enzyme expression rate *H_z_* physiologically permitted. Regulation is then designed to ensure that enzyme production is shut down after only a short burst of expression, which often also involves high levels of repressor expression (Fig. 3BEH). However, as long as there is a cost for expression of repressor, a local optimum can exist at an intermediate level of enzyme expression rate (Fig. 3GH). From here on, we will consider the maximal rate of enzyme expression *H_z_* fixed at a high level and focus on the design of regulation that optimizes the response accordingly.

**FIG. 3:**
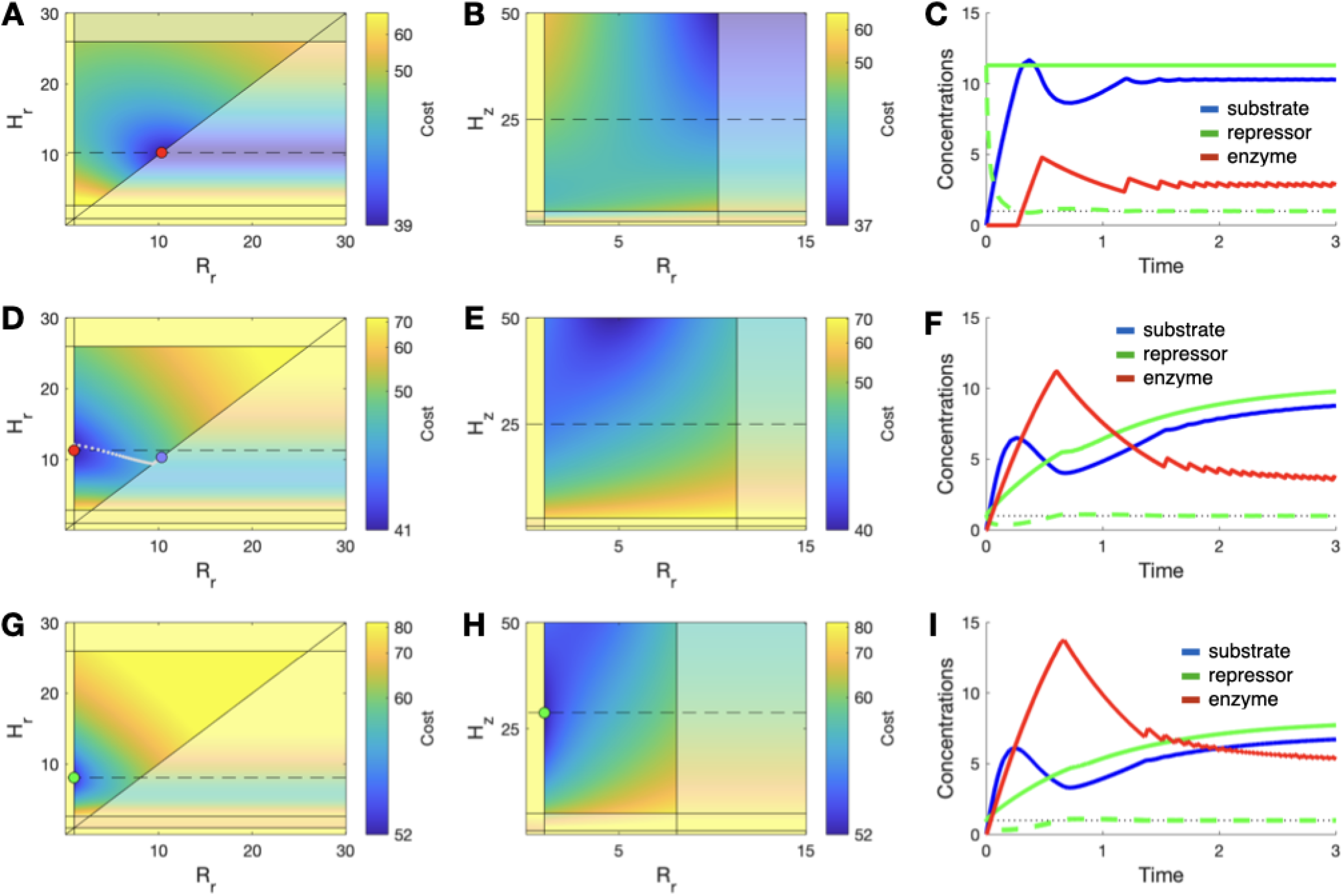
Selective pressures to minimize enzyme or substrate concentration result in qualitatively different optimal regulatory strategies. **A** Cost of the response across the *R_r_* × *H_r_* plane for *K_i_* = 1, *H_z_* =25, *S* = 50 and weights [*w_ri_ w_zi_ w_zm_ w_sm_ w_re_ w_ze_ w_se_*] = [0 1021011], where peak enzyme concentration is relatively more costly for the cell than peak substrate concentration. The cost of expressing enzyme in the absence of substrate is very high, but there is no cost of regulation. Optimal parameters are indicated by the red dot, located at the edge corresponding to a constitutively expressed repressor. **B** Cost of the response across the *R_r_ × H_z_* plane for the same parameters as A. *H_r_* is at the optimal value calculated in A, and the dashed line corresponds to *H_z_* = 25 used in A. **C** Time course of the response for the optimal parameters calculated in A, with low peak concentration of enzyme. **D-F** Cost and time course of the response for the same parameters as in A, except for weights *[w_ri_ w_zi_ w_zm_ w_sm_ w_re_ w_ze_ w_se_*] = [0 10 1 2 0 1 1], where peak substrate concentration is more costly for the cell than peak enzyme concentration. The optimum is now located at the edge corresponding to a strongly self-repressed repressor (red dot), and the time course of the response shows a low peak substrate concentration. Intermediate optima found when weights shift linearly from values in A (blue dot) to values in B are shown in gray. **G-I** Cost and time course of the response for the same parameters as A, except for weights [*w_ri_ w_zi_ w_zm_ w_sm_ w_re_ w_ze_ w_se_*] = [11012111], when there is a cost of regulation. The optimum (green dot) is now found for an intermediate value of *H_z_*.

Depending on the relative costs of substrate and enzyme, the system is optimally regulated by qualitatively different strategies. Responses that prioritize minimization of enzyme levels show optimal responses with constitutive repressor expression, while responses that prioritize minimization of substrate levels require strong self-repression of the repressor (Fig. 3A-F). To avoid any unnecessary expression of enzyme, the cell must keep high concentrations of repressors throughout the response. Therefore, selective pressures with a high cost of peak enzyme concentration find optima along the *R_r_* = *H_r_* boundary, with high constitutive expression of the repressor. However, this solution comes at the expense of a slow induction of the expression of the enzyme, since intracellular substrate levels need to rise further to bind and deplete a large initial pool of repressor. To avoid the resulting accumulation of substrate the response must follow the inverse logic. For a rapid response, where the enzyme is induced quickly upon exposure to the substrate, the initial concentration of repressor must be as low as possible. Therefore, selective pressures with a high cost of the maximal substrate concentration find optima near the *R_r_* = 1 boundary (*R_r_* = *R_z_*, in dimensional units), where the threshold for repressor self-repression is just above the threshold for enzyme repression. This solution requires the strongest possible repressor self-repression while still being able to shut off the enzyme in the absence of substrate.

## SET OF PARETO-OPTIMAL SOLUTIONS SHOWS DISTINCT STRATEGIES OF REGULATION

In order to determine the set of all possible optimized responses, regardless of selective pressures, we numerically determined the parameter sets corresponding to Pareto-optimal regimes [19–21]. Phenotypes that perform best at one single metric while disregarding all other metrics are called archetypes, and often involve extreme solutions (Fig. 4A). A phenotype involving multiple metrics is said to be Pareto optimal if, due to constraints inherent to the system, it is impossible to find any other phenotype where all metrics are simultaneously improved. For a Pareto-optimal phenotype, improving one metric necessarily decreases the performance of at least one other metric. The evolution of a given phenotype will try to improve a certain combination of the metrics that reflects the selective pressures applied to the system. Since improving all metrics together is always desirable, evolution towards Pareto-optimal solutions does not depend on the selective pressures applied. Therefore, evolution of responses under any selective pressure should bring the system to an optimal response belonging to the Pareto front, which contains all Pareto-optimal solutions. To identify the Pareto front in our system, we compute all metrics for each point in the parameter space, and then numerically eliminate points for which all metrics can be simultaneously improved [22] (so-called dominated solutions, SI). In agreement with our arguments above, the Pareto front containing all optimal solutions for any possible selective pressure applied to the system coincides with region 4 of the parameter plane (Fig. 4B).

**FIG. 4:**
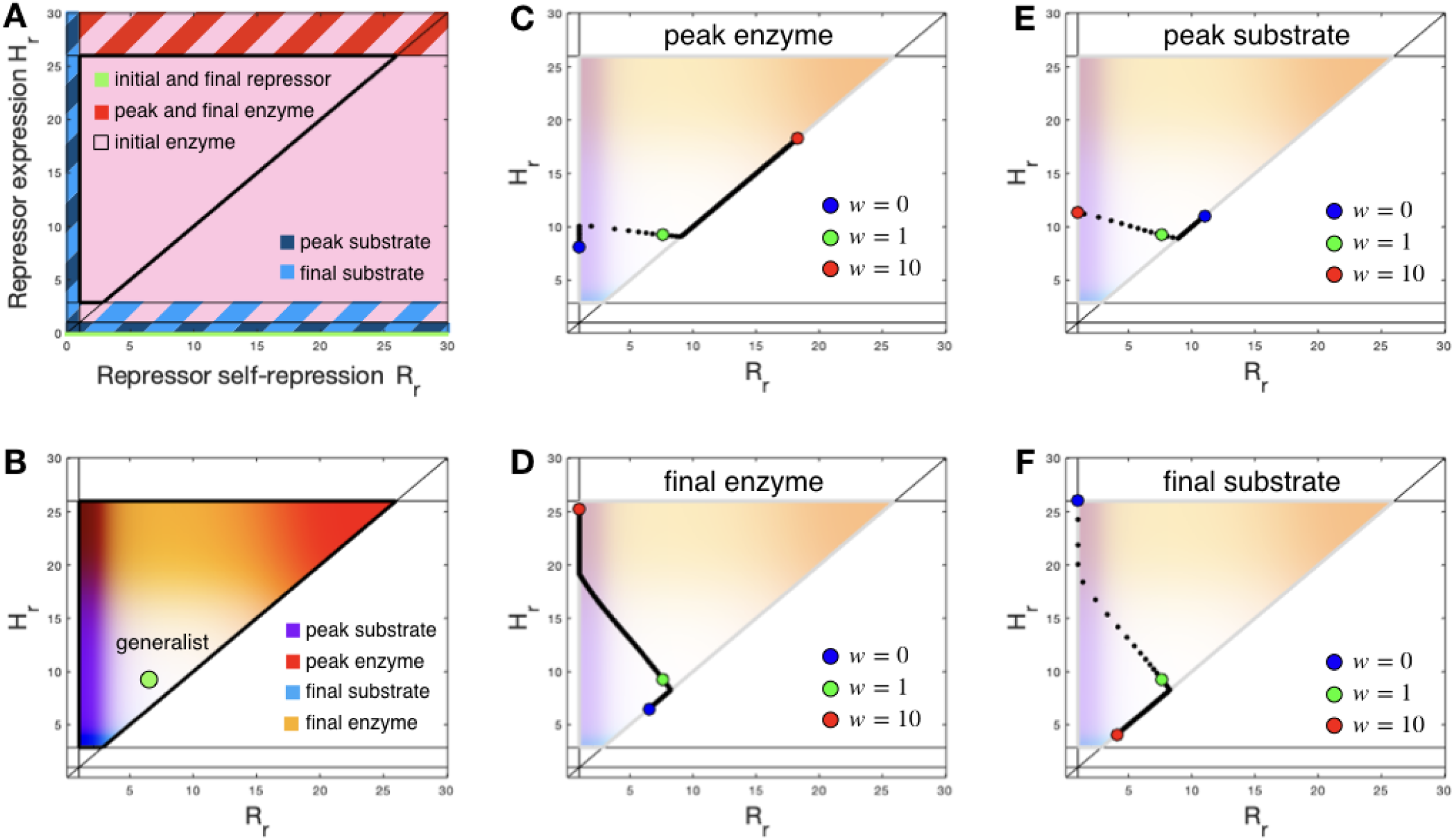
Changes in relative weight of selective pressures cause optimal regulatory strategy to switch. **A** Archetypes minimize each selective pressure while disregarding the others, leading to extreme values mostly outside of the region corresponding to regulated responses. **B** The parameter region corresponding to regulated responses is the Pareto front of optimal solutions. The edges of this region each minimize one of the selective pressures of peak and final enzyme and substrate concentrations, as indicated by the colors. The green dot in the middle of the region indicates the “generalist” strategy which optimizes the response when all four indicated selective pressures have the same weight ([*w_zm_ w_sm_ w_ze_ w_se_*] = [1 1 1 1] while *w_zi_* C> 1 and all *w_r_* = 0). **C-F** Tracking of optimal solutions when the weight of each selective pressure changes linearly from 0 to 10, with all other weights held constant at 1. 200 points are shown. In each case, the regulatory strategy switches between a constitutively expressed and a self-regulated repressor, passing through the generalist strategy.

The Pareto set of optimal solutions shows general design principles of gene regulation. We calculated how each metric (key concentrations) varies across the set, showing the trade-offs in performance among the Pareto-optimal solutions. A “generalist” response, which balances minimizing both maximum and final concentrations of both substrate and enzyme (equal weights for *s_m_, s_e_, z_m_* and *z_e_*), is optimized by parameters in the middle of the Pareto set, with a moderately expressed and weakly self-regulated repressor. However, we find that parameter sets that prioritize single traits are located along the boundaries of the region. The strength of repressor expression Hr can be adjusted to control final levels of enzyme and substrate, with weak repressor expression resulting in minimal levels of substrate and strong repressor expression resulting in minimal levels of enzyme. Conversely, the threshold of repressor self-repression *R_r_* can be adjusted to optimize maximum levels of enzyme and substrate, with strong self-repression (*R_r_* ≈ 1) resulting in minimal peak levels of substrate and weak self-repression (constitutive repressor expression, *R_r_* ≈ *H_r_*) resulting in minimal peak levels of enzyme (Fig. 4B).

Although generalist responses may be optimized by a weakly self-repressed repressor, most responses are optimized by either constitutive or strongly self-repressed repressors. Analyzing the locations of optima within the Pareto front for different combinations of weights, we note that they typically occur along the edges: either at the line *R_r_* = 1 (strong self-repression) or along the diagonal *R_r_* = *H_r_* (constitutive repressor expression). Varying the weight of each of the metrics *s_m_, s_e_, z_m_* and *z_e_*, we find that the optimum starts by moving along one edge, then quickly transitions through the generalist optimum towards the opposite edge. (Fig. 4C-F). Optimal parameters in the interior of the Pareto front, while possible, require careful balancing of the weights for all metrics and appear to be much less typical.

## DISCUSSION

Although it is not always possible to determine exactly which selective pressures result in a given set of optimized regulation parameters, the choice of regulatory strategy still reflects which costs are most critical in the evolution of a response. In some cases where not all of the key concentrations contribute towards the cost of the response, the enzyme can be regulated (or not) as one the archetypes. The gene *ampC*, which produces an enzyme that confers resistance against *β*-lactam antibiotics, is notably expressed constitutively in low levels in *E. coli*, suggesting that at typical drug concentrations for this organism the level of enzyme necessary to inactivate the drug does not pose significant costs for the cell. However, many pathogenic strains of *E. coli* that are exposed to higher antibiotic concentrations develop mutations that increase expression of AmpC, which incurs fitness costs for the cell [23, 24]. In typical responses where multiple costs are relevant, however, responses are regulated. Therefore, in many pathogen species *ampC* is expressed much more strongly, but regulated by a self-regulated repressor *ampR*, which indicates that the cost of expressing the enzyme is relevant for these organisms [25, 26]. Conversely, repressor LexA of the SOS response is often lost in pathogenic spirochetes when these species are continuously exposed to DNA damage, which results in a constitutively expressed response [27, 28]. We next characterize the regulatory strategies of several types of cell responses that are well described in the literature by calculating the regulatory parameters from data on protein expression and placing them into our optimization framework (Fig. 5, details in the SI).

**FIG. 5:**
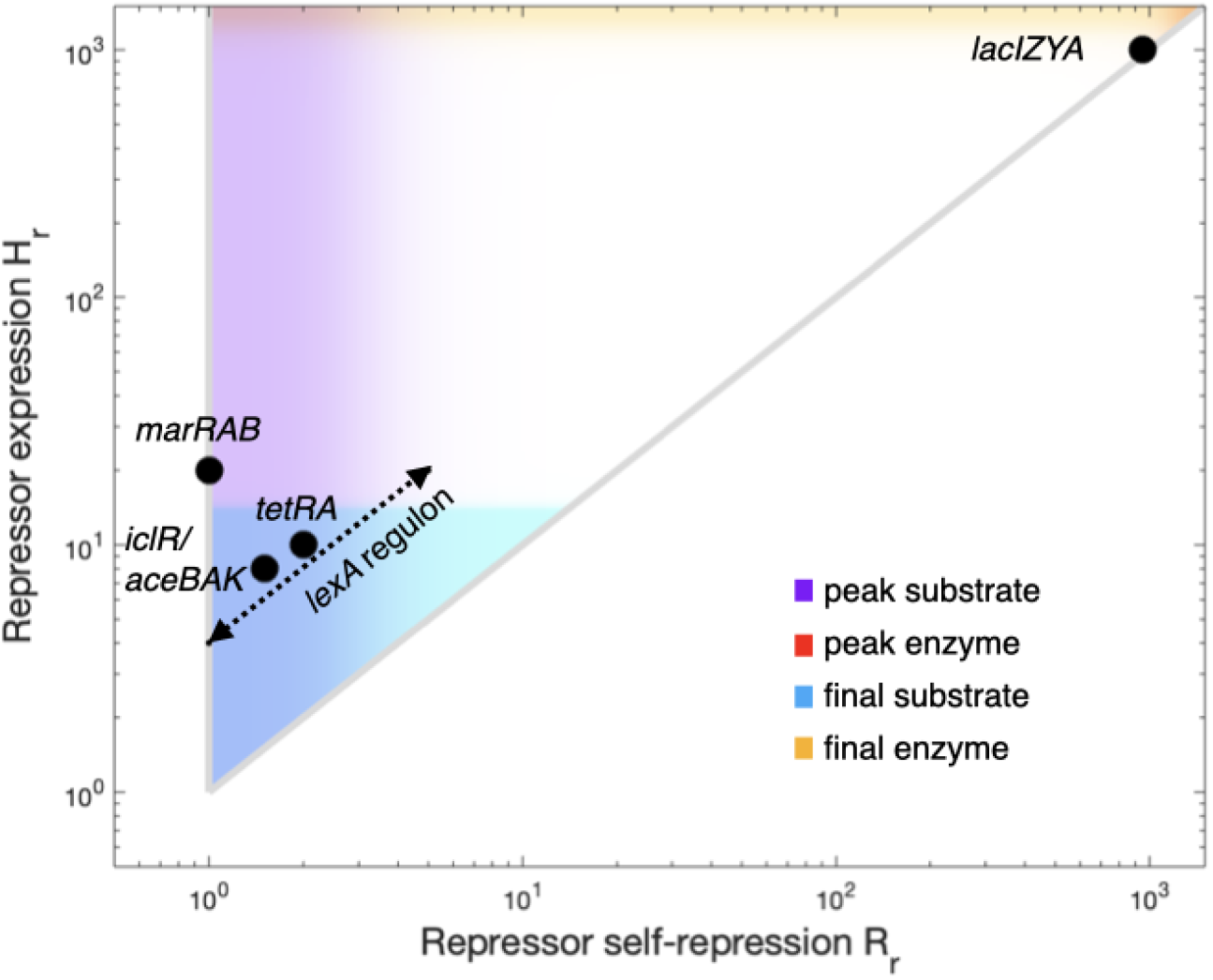
Natural examples of cellular response systems show different strategies of regulation. Regulatory parameters were calculated for a variety of cell responses. Antibiotic resistance mechanisms, such as the tetracycline resistance *tet* operon repressed by TetR and the multidrug resistance *marRAB* operon repressed by MarR, present strong repressor selfrepression and moderate repressor expression. The lactose utilization *lac* operon is repressed by a constitutive repressor LacI, expressed much above the threshold for repression of the *lac* genes. The acetate utilization *aceBAK* operon is repressed by a strongly self-repressed IclR repressor, as in antibiotic responses. The SOS response to DNA damage involves the coordination of many genes regulated by the LexA repressor, which are expressed by promoters with different affinities to LexA to ensure expression in a temporally structured fashion. Different genes in the LexA regulon are therefore located in different positions along the dotted line. The edges of the Pareto front that minimize each of the selective pressures of peak and final enzyme and substrate are indicated by the colors described in the legend.

Antibiotic responses are very often regulated by strongly self-repressed repressors, which minimizes the peak antibiotic concentration reached during the response. Negative auto-regulation has been shown to significantly increase the speed of transcriptional responses [29, 30], which is essential to keep peak drug levels low. Negative auto-regulation also decreases noise and increases the input dynamic-range of responses, which makes the system more adaptable to different drug concentrations [31–33]. In the tetracycline resistance *tet* operon, repressor TetR is expressed from a promoter that is marginally less repressed than the tightly controlled promoter that expresses efflux pump TetA [34]. Therefore, when TetR levels drop slightly, the repressor pool is replenished before the costly induction of the efflux pump, protecting the system against unnecessary activation [7].

The *marRAB* operon is induced by phenolic compounds and activates expression of global activator MarA, which is not an enzyme that acts directly on its inducers, but instead initiates a broad response that controls a wide array of genes that improves resistance against many stressors, including multiple antibiotics [35, 36]. The repressor of the operon, MarR, is expressed from the same promoter as MarA, so when MarR levels drop slightly MarA is also expressed. However, full expression of the operon depends also on a positive feedback loop of MarA activation, which occurs only at higher expression levels [37–39]. This mechanism also prevents unnecessary activation of the response. Parameter Hr, which measures expression of repressor in relation to its ability to repress enzyme, is typically not very high in antibiotic responses, since that would increase steady-state levels of drug inside the cell.

Stress responses also follow the same pattern of strongly self-repressed repressors with a relatively low *H_r_*. The SOS response to DNA damage is induced by single stranded DNA, which causes cleavage of global repressor LexA, resulting in the activation of dozens of genes involved in DNA repair [40]. Similarly to the *marRAB* operon, the DNA repair enzymes activated in the SOS response do not act directly on the inducers of the response, but they do indirectly cause their concentration to fall. This response requires coordinated expression of genes in the LexA regulon, so promoters with different affinity for LexA are used to control repair genes according to their temporal sequence. While some genes have lower thresholds of repression, as *lexA* itself, and are induced immediately as LexA levels begin to drop, other genes are very tightly repressed, and are only expressed later in the response [41].

Metabolic responses are often regulated by constitutively expressed repressors, which minimizes peak enzyme levels at the expense of a slower response and a higher peak of substrate concentration [42]. Such is the case of the *lac* operon, which produces enzymes LacZ, LacY and LacA necessary for lactose metabolism. These enzymes are repressed by repressor LacI, which is expressed nearly constitutively [43]. Although failure to quickly process lactose might result in a competitive disadvantage for the cell, high intracellular concentration of a nutrient such as lactose is not costly per se, as it is not toxic. Therefore, unlike the case of antibiotic resistance, induction of the response is less urgent. Consequently, in metabolic responses such as lactose utilization the cell prioritizes controlling enzyme levels over substrate levels, adopting a regulatory strategy that minimizes both peak and steady-state levels of the enzyme [44].

In the transition to stationary phase, *E. coli* cells induce a metabolic response that allows the consumption of the acetate secreted as a by-product during exponential growth. This response is regulated by repressor IclR which represses the *aceBAK* operon necessary for growth on acetate. Unlike LacI, IclR strongly represses its own expression [45, 46]. In this case, both the toxicity of high acetate levels and the scarcity of alternative carbon sources typical of the stationary phase increase the costs associated with substrate levels, so the acetate metabolic response has similar regulation to antibiotic or stress responses.

Although a “generalist” regulation strategy, with relatively weak negative auto-regulation of the repressor, could in theory optimize the system in situations where all costs are equally important, it is rarely observed in nature. If the generalist strategy were to be adopted, moderate changes in the relative weights of the different costs would either prompt the cell to adopt stronger self-repression of the repressor or to get rid of self-regulation altogether. Moreover, self-repression or constitutive expression of the repressor does not simply depend on minor adjustments in parameters of the regulation, but on the arrangement of regulatory elements such as promoters and repressor binding sites. Therefore, strong differences in selective pressures would be necessary to switch from one strategy to the other.

We note that in metabolic responses, the benefit of processing substrate saturates when the cell is no longer capable of utilizing the resulting metabolites. Therefore, the corresponding opportunity cost associated with unprocessed intracellular substrate saturates at higher concentrations, and does not increase in a similar fashion as the costs of intracellular stressors. Since in this case the costs associated with substrate levels are fundamentally different than the costs associated with enzyme levels, the optimal regulation might deviate from the one suggested by the analysis presented here. Similarly, with regulatory designs involving additional feedback loops can show different dynamics, as is the case with the *marRAB* operon. This is also the case for responses where the proteins under direct control of the transcription factor do not act on the substrate (or an inducer molecule) directly, so substrate levels only decrease later after the activation of enzymes further downstream. Additionally, there are plausible scenarios where the approximations used in our model would not be valid, such as when the intracellular substrate concentration is low enough to be comparable to the repressor concentration or high enough to saturate the enzyme. However, this model should hold in most cases during the initial phase of a response to a sharp increase in intracellular substrate concentration.

While we focused our analysis on the case of an enzyme directly regulated by a transcription repressor, this framework can be used to analyze other types of regulatory architectures. These include responses regulated by transcription activators or by multiple regulators, responses involving multiple enzymes or responses with complex regulation where the binding of transcription factor to DNA cannot be well approximated by “on/off” states. While an analytical solution to the model might be challenging, most of these cases could still be described by a manageable number of parameters to permit numerical solutions. Therefore, this general framework can be used to study and categorize the vast collection of regulatory architectures found in natural systems.

